# Inversion of pheromone preference optimizes foraging in *C. elegans*

**DOI:** 10.1101/2020.05.18.101352

**Authors:** M. Dal Bello, A. Pérez-Escudero, F. C. Schroeder, J. Gore

## Abstract

Foraging animals have to locate food sources that are usually patchily distributed and subject to competition. Deciding when to leave a food patch is challenging and requires the animal to integrate information about food availability with cues signaling the presence of other individuals (e.g. pheromones). To study how social information transmitted via pheromones can aid foraging decisions, we investigated the behavioral responses of the model nematode *Caenorhabditis elegans* to food depletion and pheromone accumulation in food patches. We experimentally show that animals consuming a food patch leave it at different times and that the leaving time affects the animal preference for its pheromones. In particular, worms leaving early are attracted to their pheromones, while worms leaving later are repelled by them. We further demonstrate that the inversion from attraction to repulsion depends on associative learning and, by implementing a simple model, we highlight that it is an adaptive solution to optimize food intake during foraging.

## Introduction

Decisions related to foraging for food are among the most critical for an animal’s survival. They can also be among the most challenging, because food is usually patchily distributed in space and time, and other individuals are attempting to find and consume the same resources (Abu Baker and Brown, 2014; Driessen and Bernstein, 1999; Stephens and Krebs, 1987).

An important decision, which has been the focus of considerable effort in models of foraging behavior, is for how long to exploit a food patch. Most models involve patch assessment by individuals and postulate that the leaving time depends on local estimates of foraging success (Eric L. Charnov, 1976; Oaten, 1977; Stephens and Krebs, 1987). As such, foragers are predicted to depart from a food patch when the instantaneous intake rate drops below the average intake rate expected from the environment, a phenomenon that has been observed in several animals, from insects (Wajnberg et al., 2008) to birds (Cowie, 1977; Krebs et al., 1974) and large mammals (Searle et al., 2005). The presence of other animals, however, affects individual foraging success so that different leaving times can be expected (Aubert-Kato et al., 2015; Giraldeau and Caraco, 2000). Once an animal leaves a food patch, another crucial decision is how to explore the environment to locate new sources of food. Since natural habitats are usually saturated with many different non-specific chemical cues, animals use pheromones and other odors to orientate their searches (Wyatt, 2014). This, however, implies determining whether pheromones point towards a resource supporting growth and reproduction or an already exploited one.

To acquire this knowledge, animals have to learn from experience. In the context of social foraging it has been shown that individuals might need to rely only on the most recent experience (Krebs and Inman, 1992). As such, the valence (positive or negative signal) of pheromones acquired during the most recent feeding activity is crucial for the success of the foraging process. While this has been shown in bumblebees feeding on transient resources (Ayasse and Jarau, 2014), we still don’t know whether it is important for other animals feeding in groups. Moreover, it is not clear if the ability to use associative learning (Ardiel and Rankin, 2010) to change the valence of pheromones could improve foraging success.

The nematode *Caenorhabditis elegans* is a powerful model system to investigate how information about food availability and pheromones can shape foraging in patchy habitats. *C. elegans*, indeed, feeds in large groups on ephemeral bacterial patches growing on decomposing plant material, a habitat that can be mimicked in a petri dish (Frézal and Félix, 2015; Schulenburg and Félix, 2017). Importantly, *C. elegans* can evaluate population density inside food patches using a suite of pheromones, belonging to the family of ascarosides, which are continuously excreted by worms (Greene et al., 2016; Ludewig and Schroeder, 2013). Finally, it has been shown that pheromones and food availability control the leaving times of foraging worms. In particular, the rate at which individuals abandon the patch increases when food becomes scarce and pheromones are at high concentrations (Fig. 1A) (Harvey, 2009; Milward et al., 2011).

**Figure 1.**
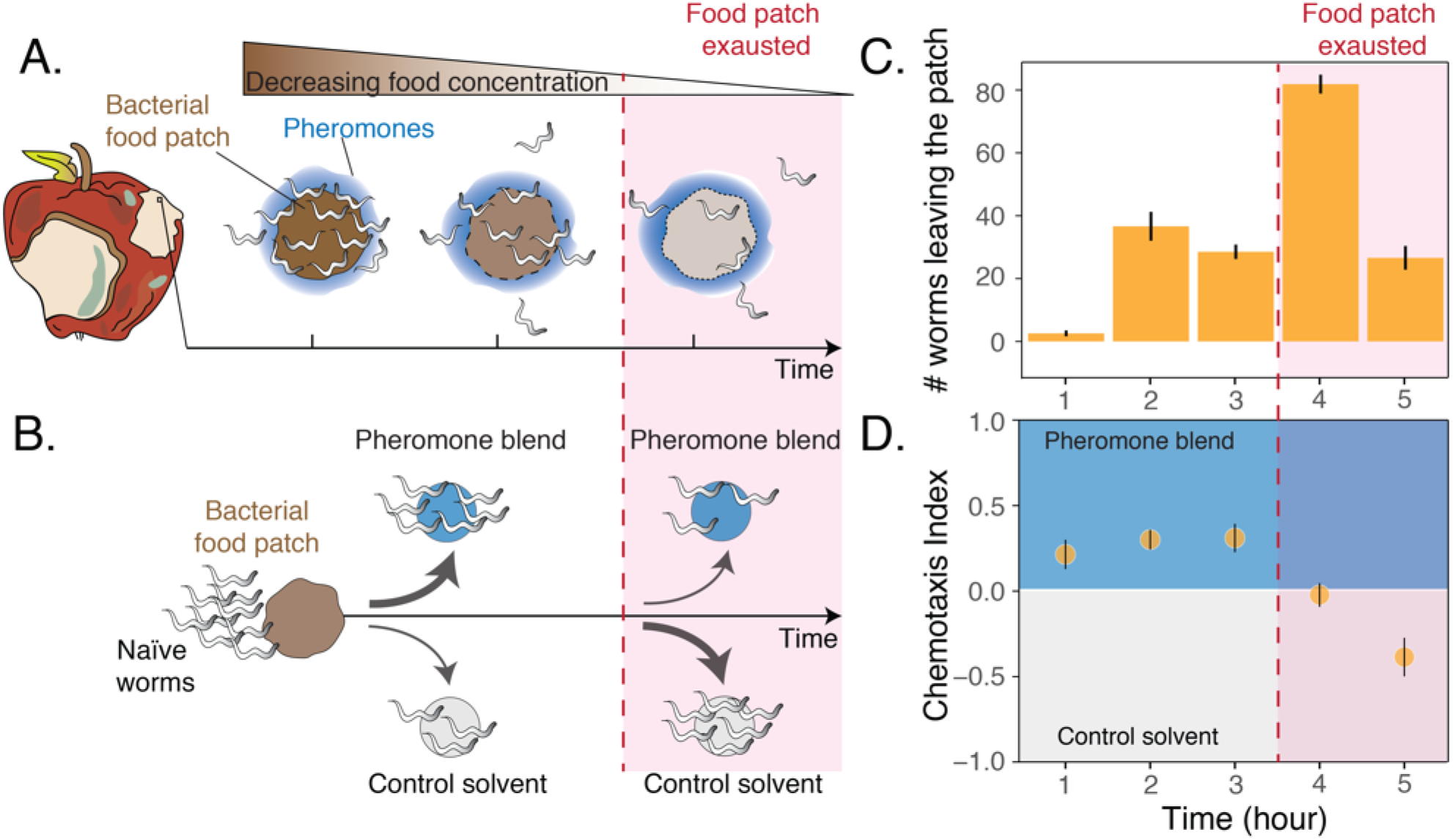
Worms leaving at different times from a food patch exhibit opposite preferences for pheromones. **A.** During the feeding process, worms remaining in the food patch experience different environmental conditions. At the beginning, food is still abundant and pheromones have already accumulated. By the end, food is scarce and pheromone concentration is even higher. **B.** In the behavioral assay, as animals feed and leave from a food patch, they are presented with the choice between a spot containing the pheromone blend and a spot containing a control solvent. In the two spots sodium azide is added in order to anesthetize the animals and prevent them from leaving the chosen spot. **C.** Individual worms leave the food patch at different times. The average number of worms that abandoned the food patch at each hour is shown (mean worm count ± SEM across replicates, n. experiments = 2). **D.** Animals leaving the food patch earlier prefer the pheromone blend while those leaving later, when the food is almost depleted, avoid the pheromone blend. In the plot, chemotaxis index is calculated on the number of naïve MY1 young adult hermaphrodites that, at each hour, reach the two spots (mean CI ± SEM across replicates, n. experiments = 2, 12 replicates each). The red region in each plot approximately indicates when food in the patch is exhausted.

In the present study, we experimentally investigated the behavioral responses of *C. elegans* to food depletion and pheromone accumulation in food patches. We confirmed that individual worms consuming a food patch leave at different times, and we found that worms leaving early have a preference for worm-secreted pheromones while those leaving late avoid the pheromones. A simple foraging model suggests that these two behaviors may optimize foraging success in the presence of competitors. Finally, using a series of behavioral assays altering worm exposure to food and pheromones, we demonstrate that associative learning underpins the change in pheromone preference.

## Results

To investigate *C. elegans* behavioral responses to food depletion, we developed an assay to simultaneously assess patch-leaving behavior and pheromone preference over time (see “Choice after food” assay in the Materials and Methods section and Fig. 1B). In our assay we used a small patch of bacteria that over the course of 5 hours is gradually depleted by feeding animals (young adult hermaphrodites of the natural isolate MY1). At equal distances from the food patch there are two spots, one of which contains a pheromone blend. Shortly after worms leave the food patch, they make a decision by choosing between the two spots (Fig. S1A). This assay mimics a foraging decision in which animals must decide when to leave a food patch and after doing so whether to follow a pheromone cue, which is indicative of the presence of others. In these experiments, the pheromone blend is obtained by collecting and filtering the supernatant of well-fed worms maintained in a liquid culture (Choe et al., 2012; Harvey, 2009; White et al., 2007). In agreement with previous results (Milward et al. 2011), we found that feeding animals abandon the food patch at very different times, with some worms leaving at the beginning while others stay until the food patch is depleted (Fig 1C). In addition, the leaving time affects worms’ preference for their pheromones, with individuals leaving the food patch early (first three hours) going to the pheromone blend (positive chemotaxis index, Fig. 1D) while worms leaving the food patch later avoiding it (negative chemotaxis index, Fig. 1D). *C. elegans* responses to food depletion, therefore, include an inversion in pheromone preference dependent on the leaving time from the food patch.

Inside rotting fruits and stems where *C. elegans* forage, bacterial food is patchily distributed (Frézal and Félix, 2015; Schulenburg and Félix, 2017). We might then expect that the timing of dispersal from existing food patches and the strategies that optimize food intake are crucial for worm survival. A natural question then arises: can the behaviors we observed in our experiments provide any benefit to *C. elegans* foraging? We explore this question with a simple theoretical model that allowed us to explore the optimality of the inversion in pheromone preference in the context of foraging in a heterogeneous environment.

The scenario described in our model implies that there are two types of food patches, those that are occupied by worms and those that are not. We assume that the occupied patches are easier to find as a result of either being nearby or because of the accumulation of pheromones secreted by the worms. Dispersing away from the occupied food patches in search for unoccupied ones therefore gives a low average payoff (*g*_*D*_), not being advantageous until the occupied food patches are depleted (see Supplement). However, the occupied patches will generically be occupied in a non-equal manner, meaning that one patch will be consumed before the other. Under these conditions, we then ask which strategy maximizes individual food intake. For the sake of illustration, In Fig. 2 we depict the simplest scenario, with two patches that are colonized by worms and another that is uncolonized.

**Figure 2.**
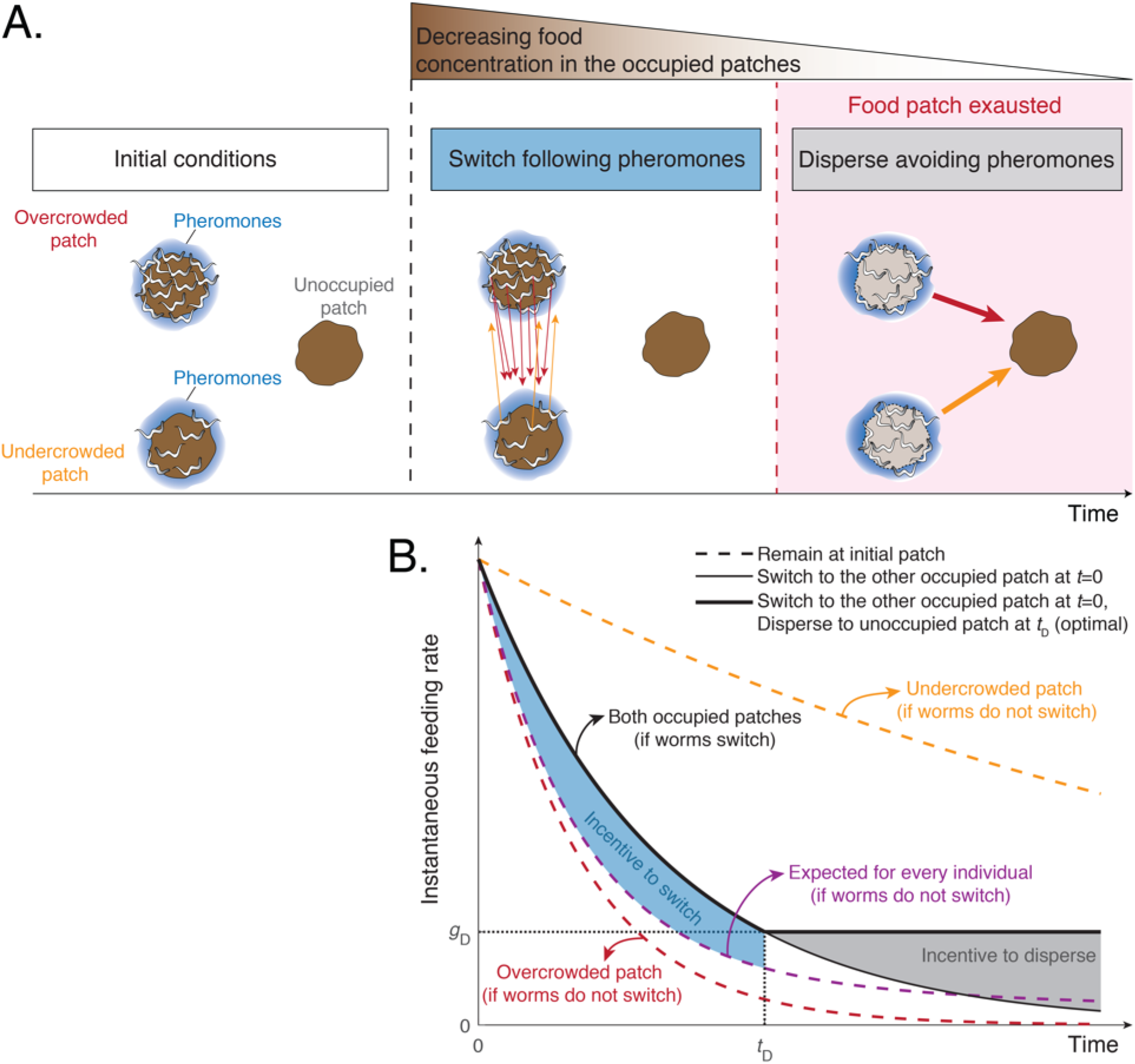
A simple model predicts that a change in pheromones preference in worms leaving late from a food patch optimizes foraging. **A.** Schematic of model predictions (only three identical food patches are depicted). Initially, two patches are unequally populated by *C. elegans* individuals (overcrowded and undercrowded food patch) while a third one is unoccupied. The release of pheromones by worms makes the occupied patches easier to find compared to the unoccupied one. During the first phase, worms equalize occupancy the occupied patches. Then, all worms stay in their patches until food becomes scarce. In this last phase, worms benefit from dispersing to the unoccupied patch avoiding pheromone cues. This would be favored by a change in pheromone preference. **B.** Instantaneous feeding rates for the scenario in panel A. Dashed lines indicate the feeding rate in case each individual remains in its original patch (red the overcrowded one and orange the undercrowded one). Solid lines indicate the feeding rate of worms that switched to the other occupied food patch following pheromones, with the thickest line indicating when this is coupled with dispersing to the unoccupied food patch by avoiding pheromones (optimal strategy).

Initially, worms should stay in the occupied patches, but they can either remain in their initial one or switch to the other occupied patch (by leaving their current patch and following pheromones to the neighboring patch) (Fig. 2A). If all worms remain in their initial patch the overcrowded food patch will be depleted faster, so worms occupying it will benefit from switching to another occupied patch (Fig. 2B). Worms have no way to tell whether they are in the overcrowded or in the undercrowded patch, but since the majority of individuals are in the overcrowded one, every individual has a higher probability of being there. Therefore, all worms have an incentive to switch to the other occupied patch during the initial phase (blue area in Fig. 2B).

How many worms should then switch? If we assume that worms attempting to switch may in fact end up in any of the occupied patches (including the initial one) with equal probability, then all worms should attempt to switch. This result is independent of any other parameters, such as number of patches or initial distribution of worms (see Supplement). If we assume that switching to other occupied patches is costly, then the optimal switching probability will be lower, but the predictions of the model remain qualitatively unaltered (see Supplement). This model therefore predicts an Evolutionary Stable Strategy (that also happens to maximize food intake) in which some worms leave a patch before it is depleted and follow the pheromone cue (Fig. 2A). This initial phase helps equalize worm occupancy and feeding across food patches.

Once worm numbers are equalized in the two easy-to-find food patches, worms feed until the food becomes scarce. At this point, worms benefit from leaving the depleted patches (gray area in Fig. 2B) and avoiding pheromones, since pheromones now mark depleted food patches. The inversion of pheromone preference therefore helps worms to disperse to unoccupied food patches. This simple model thus predicts both different leaving times and the inversion of pheromone preference and highlights that, together, these phenomena might maximize food intake of worms foraging in a patchy environment.

Nevertheless, the model does not encode any specific mechanism underpinning the inversion of pheromone preference. The most parsimonious explanation is that animals leaving the patch earlier might differ from worms leaving later simply due to their feeding status. Indeed, early-leaving worms abandon the food patch when food is still abundant and therefore, they are more likely to be well-fed. By contrast, worms leaving later—when the food is scarce—are more likely to be famished.

However, worms leaving earlier are also exposed to pheromones in the presence of abundant food, while worms leaving later experience high levels of pheromones in association with scarce food. These conditions are analogous to those that have been shown to support associative learning in *C. elegans* (Ardiel and Rankin, 2010). Similarly to the well-known case of associative learning with salt (Hukema et al., 2008; Saeki et al., 2001), in our experiment worms could be initially attracted to pheromones because of the positive association with the presence of food. Attraction could later turn into repulsion if worms start associating pheromones with food scarcity.

To distinguish between the change in pheromone preference being caused by feeding status alone or by associative learning, we performed experiments in which young adult hermaphrodites were conditioned for five hours in the four scenarios corresponding to the combinations of +/− *food* and +/− *pheromone blend*. After conditioning, animals were assayed for chemotaxis to the pheromone blend (see Materials and methods section, Fig. 3A and S1B). We found that worms go to the pheromone blend when they are conditioned with *+ food + pheromone blend* whereas they avoid it when they are conditioned with *− food + pheromone blend* (Fig. 3B, blue and yellow bars, mean CI_++_ = 0.38 ± 0.07 vs mean CI_−+_ = – 0.15 ± 0.04). Interestingly, worms conditioned without the pheromone blend do not exhibit a particular preference for their pheromones (Fig. 3B, red and turquoise bars, mean chemotaxis index for the *+ food − pheromone blend* and the *− food − pheromone blend* scenarios are 0 ± 0.07 and – 0.02 ± 0.06, respectively). Worms therefore exhibit attraction when pheromones are paired with abundant food and aversion when pheromones are associated with absence of food. Otherwise, *C. elegans* does not show a specific preference for pheromones. These findings are consistent with the hypothesis that the *C. elegans* preference for the pheromone blend changes due to associative learning.

**Figure 3.**
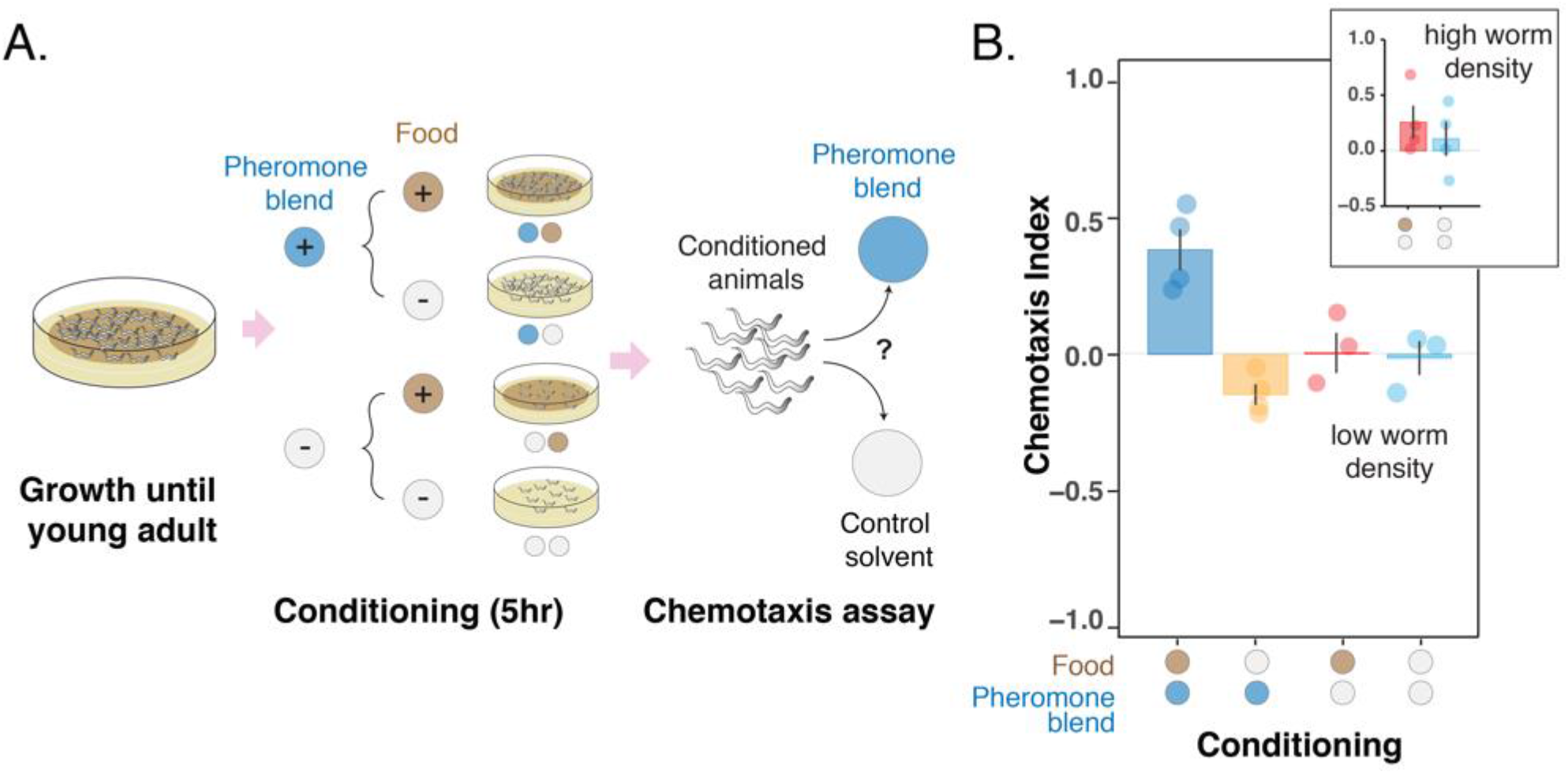
Changes in pheromone preference depend upon associative learning. **A.** MY1 individuals grow at high density and with plenty of food until young adult. Animals are then transferred to conditioning plates, where they spend 5 hours. Conditioning scenarios are four: *+ food + pheromone blend; − food + pheromone blend; + food− pheromone blend; − food − pheromone blend*. To prevent uncontrolled pheromone accumulation, the conditioning scenarios without added pheromone blend had to be repeated at low worm density. Worms are then assayed for chemotaxis to the pheromone blend. **B.** MY1 individuals are not generally attracted the pheromone blend unless it is paired with abundant food. Chemotaxis index is shown for the four different conditioning scenarios: *+ food + pheromone blend* (blue bar); *− food + pheromone blend* (yellow bar); *+ food− pheromone blend* (red bar); *− food − pheromone blend* (turquois bar). As a comparison, chemotaxis index is shown for the *+ food− pheromone blend scenario* and the *− food − pheromone blend* scenarios with conditioning done at normal animal density (Panel B, inset). Points indicate the outcome of each independent replicated experiment (n=4 and n=3 for experiments with worms at low population density) while bars indicate mean CI ± SEM across independent experiments.

Conditioning in the two scenarios without pheromone blend added had to be performed at low worm density due to uncontrolled pheromone accumulation. Indeed, when conditioned at high worm density, animals in the *+ food − pheromone blend* scenario were still exposed to the pheromones that they kept excreting during the 5-hour conditioning period (Sakai et al., 2013) and therefore exhibited attraction to the pheromone blend, albeit variable (mean CI_+−_ = 0.25 ± 0.07, Fig. 3B inset, red bar). Worms conditioned in*− food − pheromone blend* displayed no significant attraction to the pheromone (mean CI_−−_ = 0.10 ± 0.15, Fig. 3B inset, turquois bar). The variation here is even bigger, likely due to the fact that the pheromone cocktail produced by starved worms can be different from the pheromone blend we used, which was obtained from well-fed worms (Kaplan et al., 2011).

The outcome of this series of experiments with the pheromone blend can be recapitulated with pure synthetic ascarosides (Fig. S2). This allowed us to address the fact that the pheromone blend contains, in addition to a cocktail of ascarosides, other products of worm metabolism, compounds deriving from the decomposition of dead worms and bacteria, and perhaps other unknown substances. Overall, these results provide additional evidence that it is associative learning with ascaroside pheromones that underlies the inversion of pheromone preference observed in our original foraging experiment (Fig. 1).

To provide further support that the *C. elegans* preference for pheromones can change through associative learning, we asked whether the change in preference occurs also via the association with a repellent compound, namely glycerol (Hukema et al., 2008). To answer this question, we performed a learning experiment in which young adult hermaphrodites were conditioned for one hour in four different scenarios deriving from all the possible combinations of +/− *repellent* (glycerol) and +/− *pheromone blend*. During conditioning, animals are free to dwell in a plate seeded with *E. coli* OP50, and therefore they are always exposed to a high concentration of bacterial food. Here, uncontrolled pheromone accumulation was not an issue thanks to the short conditioning period. After conditioning, worms are tested for chemotaxis to the pheromone blend (Fig. 4A).

**Figure 4.**
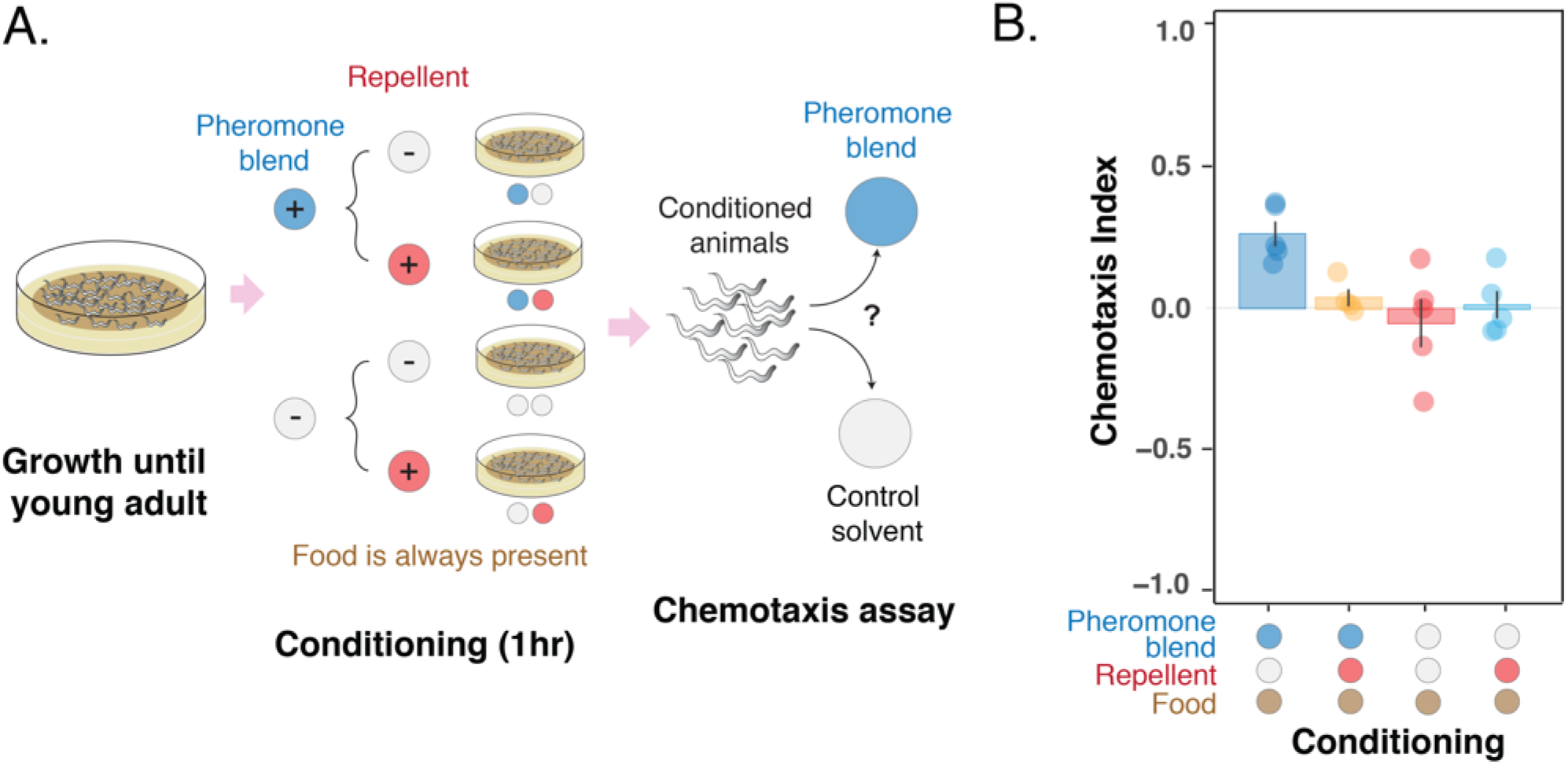
Pheromone preference changes due to association with the presence or absence of a repellent compound (glycerol). **A.** MY1 individuals grow at high density and with plenty of food until young adult. Animals are then transferred to conditioning plates, where they spend 1 hour. Conditioning scenarios are four: *+ pheromone blend − repellent; + pheromone blend + repellent; − pheromone blend − repellent; − pheromone blend + repellent*. Bacterial food is always abundant to prevent confounding effects due to the feeding status of animals. The short conditioning time prevents uncontrolled pheromone accumulation. Worms are then assayed for chemotaxis to the pheromone blend. **B.** MY1 individuals are not attracted to the pheromone blend unless it is present and paired with food. Association with the repellent disrupts worm preference for pheromones gained in the presence of abundant food. Chemotaxis index is shown for the four different conditioning scenarios: *+ pheromone blend − repellent* (blue bar); *+ pheromone blend + repellent* (yellow bar); *− pheromone blend − repellent* (red bar); *− pheromone blend + repellent* (turquois bar). Points indicate the outcome of each independent replicated experiment (n=5) while bars indicate mean CI ± SEM across independent experiments.

We found that the preference for the pheromone blend, which is retained in the *− repellent + pheromone blend* scenario (Fig. 4B, blue bar, mean CI_−+_ = 0.26 ± 0.04), is lost in the *+ repellent + pheromone blend* condition (Fig. 4B, yellow bar, mean CI _++_ = 0.04 ± 0.03). Again, when animals are conditioned in the absence of pheromones, they do not show any particular preference for the pheromone blend (mean CI_+−_ = 0.01 ± 0.05, mean CI_−−_ = − 0.05 ± 0.08, Fig. 4B red and turquois bars respectively). In other words, animals do not exhibit a particular preference for pheromones, except when they are exposed to pheromones and food (in the absence of the repellent). Exposure to the repellent in the presence of food and pheromones is required to disrupt attraction. The outcome of this experiment provides further support that *C. elegans* can change its preference for the pheromone cocktail it produces through associative learning.

## Discussion

We have assessed the response of *C. elegans* to food depletion and how this influences worms’ response to their pheromones. In agreement with previous studies, our findings indicate that worms exhibit different leaving times when feeding in groups on transient bacterial food patches. Interestingly, the leaving time affects *C. elegans* preference for its pheromones, with animals leaving early being attracted to their pheromones and worms leaving later being repelled by them. We showed that this inversion from attraction to repulsion depends on associative learning and appears to be an adaptive solution to optimize food intake during foraging.

Our model shows that a change from pheromone attraction to repulsion is required to optimize food intake when three different factors are combined. First, food patches that give diminishing returns, which should be abandoned when the environment provides a better average intake rate. This first factor has been thoroughly studied both theoretically and experimentally in the context of optimal foraging and the marginal value theorem (Eric L. Charnov, 1976; Krebs et al., 1978; Oaten, 1977; Stephens and Krebs, 1987; Watanabe et al., 2014), and in this respect our model simply reproduces previous results.

The second factor is competition for limited resources, which in our model creates the need to switch between pheromone-marked food patches in order to distribute the individuals more evenly. This redistribution closely resembles the Ideal Free Distribution, which postulates that animals should distribute across patches proportionally to the resources available at each source (Bautista et al., 1995; Fretwell and Lucas, 1969; Houston and McNamara, 1987; Kennedy and Gray, 1993). However, here, our model does depart from previous studies. The Ideal Free Distribution applies to cases in which the benefit per unit time decreases with the number of individuals exploiting the same resource. This is the case for habitat choice (where the distribution was first proposed, Fretwell and Lucas, 1969), or if the instantaneous feeding rate decreases with the number of feeders (Houston and McNamara, 1987). It is not however the case in many foraging scenarios, including the one represented by our model (and implicitly by most optimal foraging models), in which animals can feed unimpeded by each other. In these cases, a higher number of animals simply means that the resource is depleted faster (this can be seen in Fig. 2B: At time *t*=0, animals feed at the same rate in both food patches). Simply adding competition to standard optimal foraging models will not change their results qualitatively. Animals will stay in each food source until the food is so scarce that the instantaneous feeding rate falls below the environment’s average. This will happen earlier for more crowded food sources, but animals will never need to switch across food sources before they are depleted.

The third key factor in our model is non-stationarity: We assume that all pheromone-marked food patches were colonized and will be depleted at roughly the same time. This fact creates the need to switch before the current patch is depleted, because by then most of the benefit from undercrowded (but pheromone-marked) food patches will be gone. This non-stationary environment has received less attention than the previous factors. It is typical of species with boom-and-bust life cycles such as *C. elegans* (Frézal and Félix, 2015), but may also be applicable to other cases, such as migratory species (which arrive synchronously to a relatively virgin landscape), fast-dispersing invader species or, in general, species that occupy a non-stationary ecological niche.

*C. elegans* individuals use stimuli coming from the environment (smells, tastes, temperature, oxygen and carbon dioxide levels) and from other individuals (pheromones) to efficiently navigate their habitat. An important evolutionary adaptation in this regard is that the *C. elegans* preference for each of these stimuli can change through experience, including acclimation (Fenk and de Bono, 2017) and associative learning phenomena (Ardiel and Rankin, 2010; Colbert and Bargmann, 1995; Rankin, 2004). We have identified associative learning as the most plausible phenomenon underpinning the change in pheromone preference. During feeding, worms learn to give a positive or negative preference to pheromones depending on the context in which they experience them, in particular the presence or absence of food (Wyatt, 2014). A similar learning process occurs in bumblebees that, in their natural habitat, do not land or probe flowers that have been recently visited and marked by chemical footprints left by themselves or other bees. It has been shown that only experienced foragers, i.e. those that learnt to associate the chemical footprints with the absence of nectar in marked flowers, can successfully avoid them and increase their overall nectar intake (Ayasse and Jarau, 2014). This suggests that associative learning based on pairing pheromones or similar chemical signals with food availability might be frequently observed in animals feeding in groups, not only eusocial insects, as a strategy to increase food intake.

We have shown that dispersal of feeding stages of *C. elegans* from occupied patches is regulated by the recent experience of food availability and pheromones, which indicates, at any time, whether it is better to follow the scent of pheromones or to avoid it. A mechanism based on the synergistic interaction between food and pheromones also regulates *C. elegans* dispersal over longer time scales and, in general, its boom-and-bust life cycle (Edison, 2009; Frézal and Félix, 2015). Indeed, scarce food and high concentration of pheromones promote the entry into a resting stage (the dauer larva), allowing worms to survive unfavorable seasons and disperse to uncolonized rotten material, where abundant food, in turn, resumes development to adulthood (Frézal and Félix, 2015). Our findings establish an interesting parallel between mechanisms promoting dispersal over short and long temporal scales and highlight the important role that non-dauer stages play in exploiting transient bacterial patches.

As a final remark, our results suggest that *C. elegans* preference for pheromones might not be innate, as it was previously stated (Greene et al., 2016; Macosko et al., 2009; Pungaliya et al., 2009; Simon and Sternberg, 2002; Srinivasan et al., 2012, 2008) and question what it means to be a naïve worm. Worms that we call “naïve” are directly assayed for chemotaxis after being simultaneously exposed to both bacterial food and ascaroside pheromones, which are continuously excreted by the animals during their growth (Kaplan 2011). Hence, it is possible that the attraction that “naïve” worms exhibit is due to the positive association with food that they learn to make during growth on the plate.

## Acknowledgments

The authors would like to thank the members of the Gore lab for comments on the manuscript, Ying K. Zhang for assistance with the synthesis of ascarosides and Jonathan Friedman for feedback on the model. The *C. elegans* strain we used was provided by the CGC, which is funded by NIH Office of Research Infrastructure Programs (P40 OD010440). This work was supported by NIH and the Schmidt Foundation. While at MIT, APE was funded by EMBO Postdoctoral Fellowship Grant ALTF 818-2014 and Human Frontier Science Foundation Postdoctoral Fellowship Grant LT000537/2015.

## Author contributions

MDB, APE and JG conceived the study. MDB performed the experiments and analyzed the data. APE built the model. FCS contributed the synthetic pheromones. All authors wrote the manuscript.

## Material and methods

### Strains and culture conditions

We used a *Caenorhabditis elegans* strain recently isolated from the wild, MY1 (Lingen, Germany). The strain has been obtained from the Caenorhabditis Genetic Centre (CGC). Animals were grown at 21-23 °C (room temperature) on nematode growth media (NGM) plates (100 mm) seeded with 200 μl of a saturated culture of *E. coli* OP50 bacteria (Stiernagle, 2006). As for OP50 culture, a single colony was inoculated into 5ml of LB medium and grown for 24 h at 37 °C.

### Pheromones

We obtained the crude pheromone blend by growing worms in liquid culture for 9 days (at room temperature and shaking at 250 rpm) (Von Reuss et al., 2012). Individuals from one plate were washed and added to a 1-liter flask with 150ml of S-medium inoculated with concentrated *E. coli* OP50 pellet made from 1 liter of an overnight culture. Concentrated *E. coli* OP50 pellet was added any time the food supply was low, i.e. when the solution was no longer visibly cloudy (Stiernagle, 2006). The pheromone blend was then obtained by collecting the supernatant and filter-sterilizing it twice. A new pheromone blend was produced every 3 months. Pure synthetic ascarosides (ascr#5 and icas#9) were obtained from the Schroeder lab and kept at −20°C in ethanol. Each time an experiment was performed, an aqueous solution at the desired molar concentration was prepared (10 μM for ascr#5 and 10 pM for icas#9). The control solvent for the pheromone blend is S-medium, while the control solvent for the pure ascarosides is an aqueous solution with the same amount of ethanol present in the ascaroside aqueous solution (Srinivasan et al., 2012).

### Choice after food assay

It is a chemotaxis assay modified from Bargmann et al. (1991) and Saeki et al. (2001), performed on naïve worms that encounter a food patch before making the choice between the pheromone blend and the control solvent. We used 100 mm NGM plates in which we deployed 20 μl of the pheromone blend, 20 μl of control solvent and 15 μl of a diluted OP50 *E. coli* culture at equal distance from each other (Fig. S1A). In the pheromone and control spots, 2 μl of 0.5 M sodium azide was added in order to anaesthetize the animals once they reached the spots. Since the anesthetic action of sodium azide lasts for about 2 hours in this set-up, another 1μl was added two hours after the beginning of the assay in both spots. ~200 naïve animals were placed close to the patch of bacteria, so that they stop and feed in the patch before they chemotaxis towards the two cues. Worms are left to wander freely on the assay plate for 5 hours. The number of worms around the two spots was counted every hour and the chemotaxis index was calculated based on the number of new worms that reached the two spots during each hour.

### Chemotaxis assay

Chemotaxis assay has been performed in 60 mm NGM plates, in which worms are given the choice between pheromone (either 20 μl of the pheromone blend or 20 μl of a pure ascaroside in aqueous solution) and a control solvent (20 μl) (Bargmann and Horvitz, 1991; Saeki et al., 2001). The two spots are deployed ~3 cm apart from each other (Fig. S1B). Shortly before the start of the assay, 1 μl of 0.5 M sodium azide is added to both spots in order to anaesthetize the animals once they reach the spots. ~50 animals are placed equidistant from the two spots and left to wander on the assay plate for 1 hour at room temperature (Fig. S1B). The assay plates were then cooled at 4°C and the number of worms around each spot was counted using a lens. The chemotaxis index is then calculated as 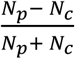, where *N*_*p*_ is the number of worms within 1 cm of the center of the pheromone spot, while *N*_*c*_ is the number of worms within 1 cm of the center of the control spot. The number of independent experiments, i.e. performed in different days, is indicated in each figure caption. For each experiment, we usually performed 10 replicated assays for each scenario.

### Learning experiment

Hermaphrodite individuals of the MY1 strain are grown until they become young adults in NGM plates seeded with 200 μl of saturated *E. coli* OP50 bacteria. Then, animals are washed off the plates with wash buffer (M9 + 0.1% triton), transferred to an Eppendorf tube and washed twice by spinning down the worms and replacing the supernatant with fresh wash buffer each time. After that, ~1000 animals are transferred to each conditioning plate. For the experiments in which worms are conditioned at low population density, the number of individuals placed in each conditioning plate is ~100. In the first series of experiments, the four different scenarios derive from all the possible combinations of +/− *food* and +/− *pheromone blend*. Plates are prepared ~16 hours before the training starts, so that bacteria can grow and form a lawn. NGM plates are seeded with +/− 200 μl of saturated *E. coli* OP50 culture and +/− 200 μl of pheromone blend. Animals spend 5 hours in the conditioning plates at room temperature before being assayed for chemotaxis to the pheromone blend. In the second series of experiments (with the repellent), the four different scenarios derive from all the possible combinations of +/− *repellent* (glycerol) and +/− *pheromone blend*. Conditioning plates are prepared ~1 before the start of the experiment and are NGM plates seeded with 200 μl of saturated *E. coli* OP50 culture +/− 0.5 M glycerol and +/− 200 μl of pheromone blend. To keep the concentration constant, when the pheromone blend was not added, we diluted OP50 with 200 μl of S-medium. Animals stay in the conditioning plates for one hour at room temperature before being assayed for chemotaxis to the pheromone blend. In the experiments with pure ascarosides *ascr#5* and *icas#9*, the four different scenarios derive from all the possible combinations of +/− *food* and +/− *pure ascaroside* (in aqueous solution) and are prepared as the experiment with food and the pheromone blend. However, the concentration of ascaroside that was added in the conditioning plate was higher than the concentration at which the worms were tested for chemotaxis (for *ascr#5* was 10 μM, while for *icas#9* was 10 pM icas#9) to compensate for the diffusion of the ascaroside throughout the agar in the conditioning plates. Ascr#5 was added at a concentration of 100 μM onto conditioning plates, while icas#9 was added at a concentration of 1 μM. Worms spend 5 hours in the conditioning plates at room temperature, after which they are assayed for pheromone chemotaxis.

## Supplementary data

**Figure S1.**
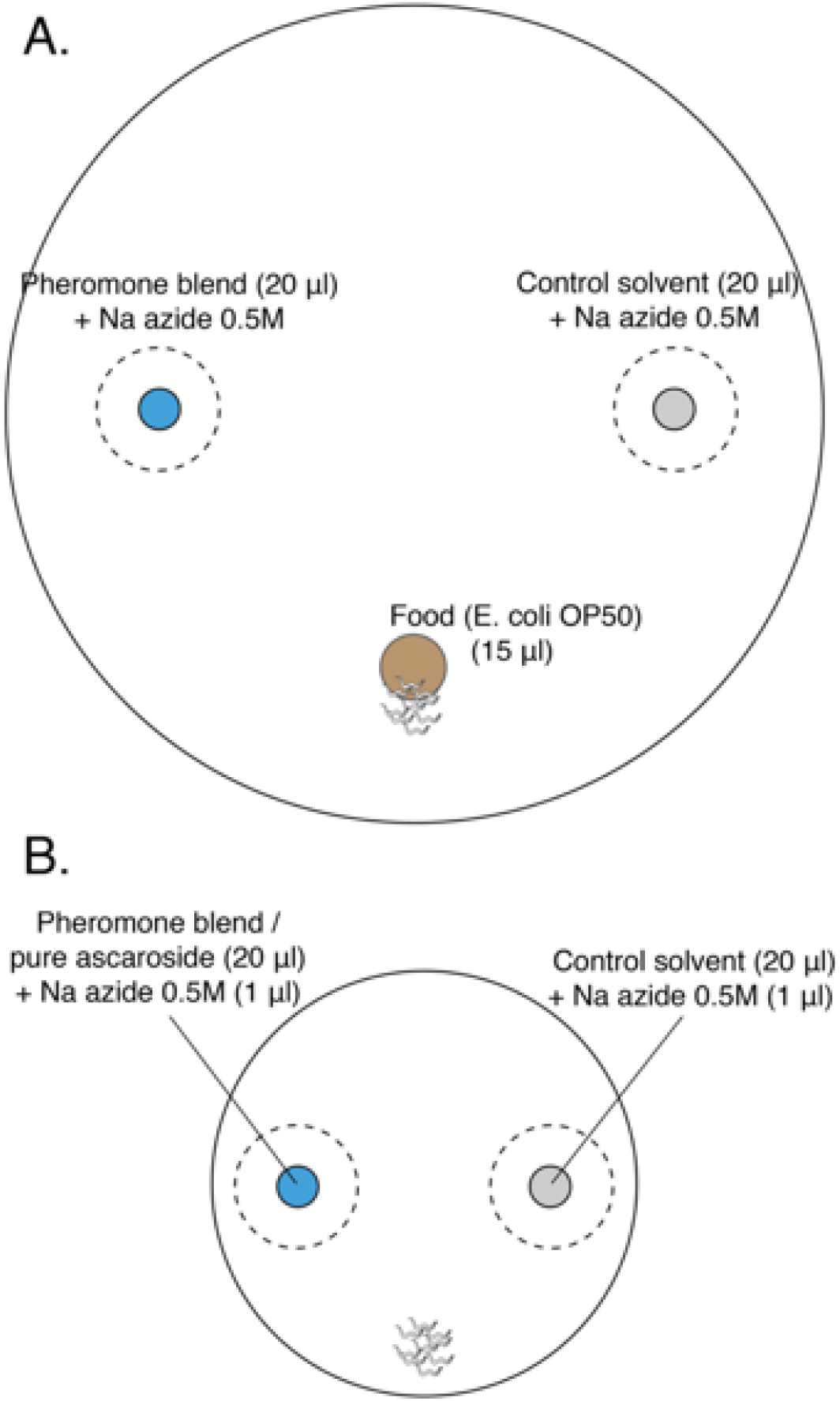
Layout of plates for chemotaxis assays. **A.** Choice after food assay: 100 mm petri dish filled with 25 ml of NGM. In blue, the spot with 20 μl of the pheromone blend, in grey the spot with 20 μl of the control solvent and in brown the food patch, 15 μl of a diluted *E. coli* OP50 culture. In the pheromone and control spots, 2 μl of 0.5 M sodium azide was added in order to anaesthetize the animals once they reached the spots. Since the anesthetic action of sodium azide lasts for about 2 hours in this set-up, another 1μl was added two hours after the beginning of the assay in both spots. **B.** Chemotaxis assay: 60 mm petri dish filled with 10 ml of NGM. In blue, the spot with 20 μl of either pheromone blend or pure synthetic ascarosides; in grey the spot with 20 μl of control solvent. In both spots, 1 μl of Na azide 0.5M is added to paralyze the worms once they reached the cue. Worms are placed equidistant from the two spots.

**Figure S2.**
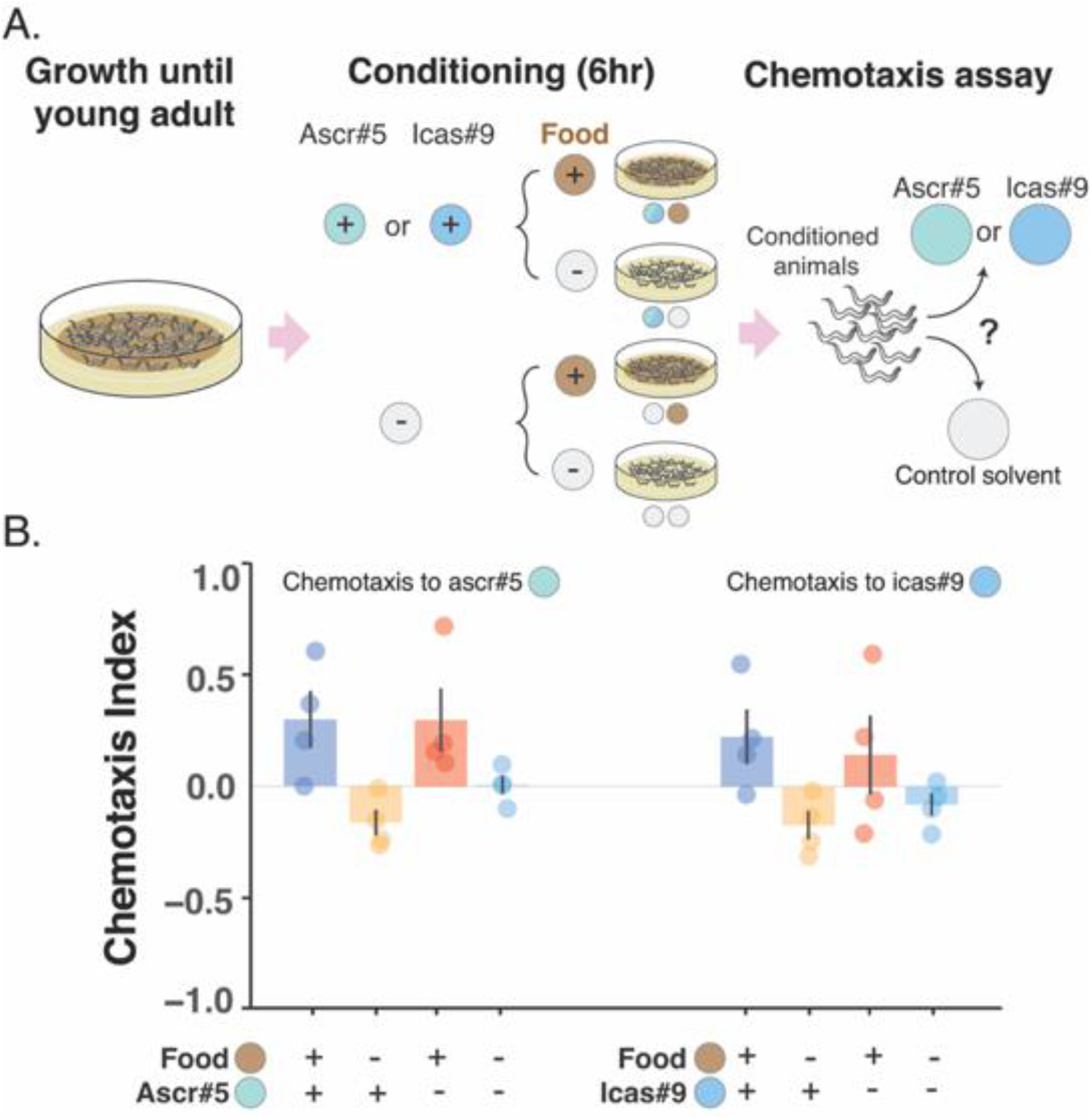
Changes in the preference for pure ascarosides, ascr#5 and icas#9, are likely to depend upon associative learning. **A.** Worms grow at high density and with plenty of food until young adult. Animals are then transferred to conditioning plates, where they spend 5 hours, before being assayed for chemotaxis to the pure ascaroside. **B.** Attraction is turned to repulsion by simultaneously pairing pure ascarosides with absence of food (associative learning). Chemotaxis index is shown for the four different conditioning scenarios: *+ food + ascaroside* (blue bars); *− food + ascaroside* (yellow bars); *+ food− ascaroside* (red bars); *− food − ascaroside* (turquois bars). Points indicate the outcome of each independent replicated experiment (n=4 for each ascaroside) while bars indicate mean CI ± SEM across independent experiments.

### Foraging model

We assume that two types of food patches exist:

- Food patches marked with pheromones, which have a high average value (they are capable of sustaining worm growth and are easy to find).
- Unmarked food patches, which have a low average value as they are more difficult to find).

Initially, individuals are distributed across the pheromone-marked patches. Let *K* be the number of patches, *n*_*i*_ the number of individuals in the *i*-th patch (for *i* = 1,2 … *K*), and *N* the total number of individuals (so 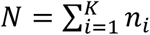).

At any time, individuals can take three possible actions: **Remain** in the current patch, **switch** to another pheromone-marked patch (so they leave the patch and follow pheromones), or **disperse** and search for an unmarked patch (so they leave the patch and avoid pheromones). Individuals’ instantaneous feeding rate *g*(*t*) depends on their choices. Let’s start with the choice of dispersal. To model this decision we borrowed the results of classical foraging models from which the Marginal Value Theorem was derived (E L Charnov, 1976). These models describe an individual depleting a food patch, whose environment contains other food patches that remain stationary (i.e. on average the other food patches are not being depleted over time). We also assume that the unmarked food patches remain stationary. In these conditions, one can compute an average expected intake rate from dispersing and searching for unmarked patches, which we will call *g*_*D*_. This average intake rate takes into account the average quality of the unmarked food patches and the time needed to find and consume them. The optimal strategy is to remain in the current food patch until the instantaneous feeding rate (*g*(*t*)) falls below *g*_*D*_ (E L Charnov, 1976). Following these models, we assume that any individual that disperses will experience a constant instantaneous feeding rate *g*_*D*_.

While we can use the formalism of classical foraging models for the dispersal decision, we cannot do the same for the switching decision, because the pheromone-marked food patches are non-stationary (they all get depleted at roughly the same time, a feature characteristic of species with a boom-and-burst life cycle such as *C. elegans*). We will therefore model explicitly food depletion in all pheromone-marked patches.

We assume that individuals at a pheromone-marked food patch feed at a rate proportional to the amount of food left in the patch: *g*(*t*) = *a*(*t*), where *a*(*t*) is the amount of food available at the food patch at time *t*.^1^ Therefore, food patches get depleted over time as *a*(*t*) = *A*_0_*e*^−*mt*^, where *A*_0_ is the initial food density in the pheromone-marked patches and *m* is the number of individuals in the patch.^2^

Individuals that switch pay a cost *c* for switching. We assume that switching is fast compared to the depletion rate of the food patches, so switching is instantaneous in our model. Individuals that switch will then arrive to any pheromone-marked food patch with equal probability (including their initial one).

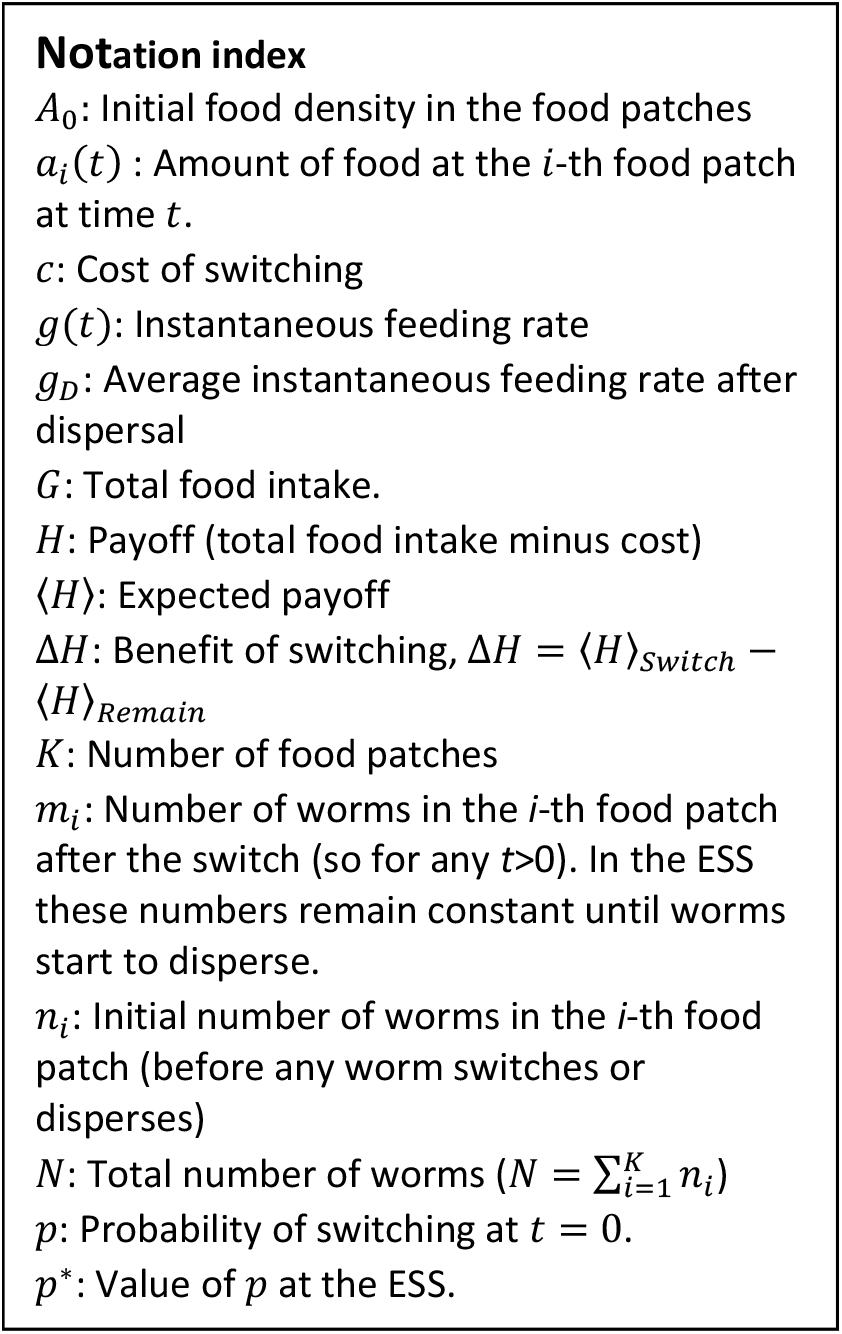

We assume that evolution has maximized the total food intake, which is the integral of the instantaneous intake rate (*g*(*t*)) over a long period of time. The exact length of this period does not actually matter, because in all relevant cases we will be comparing strategies that end with dispersal, and therefore get the same intake rate at the end. We will always work with differences between the payoffs of these strategies, so these final periods will cancel out. Therefore, for simplicity we will always integrate to *t* = ∞.

Finally, we assume that individuals have strong sensory constraints: They only perceive their instantaneous feeding rate (*g*(*t*)), not having information about any of the other parameters (number of patches, number of individuals per patch, etc.). However, their behavior can be adapted to be optimal with respect to the average values of these parameters over the species’ evolutionary history.

In these conditions, the following Evolutionary Stable Strategy (Maynard Smith, 1982, 1974) exists: At time *t* = 0, all individuals have a probability *p*\* of switching (so a fraction *p*\* of the individuals will switch). Then they all remain in the food patches until their instantaneous feeding rate falls below *g*_*D*_, at which point they disperse.

#### Proof

We will prove each part of the Evolutionary Stable Strategy separately:

##### 1. Individuals will not disperse until the food patches are depleted

Dispersing gives an instantaneous average payoff of *g*_*D*_, so individuals should never disperse if their instantaneous intake rate is above *g*_*D*_ (i.e. if the food density in their current patch is *a*(*t*) > *g*_*D*_).

##### 2. The probability to switch (*p*) has a stable equilibrium (*p*\*)

If all individuals follow the Evolutionary Stable Strategy, at time *t* = 0 a fraction *p* of them switches, changing the distribution of individuals across food patches. Let *m*_1_, *m*_2_ … *m*_*K*_ be the number of individuals in each food patch after the switch. These numbers are related to the initial distribution as

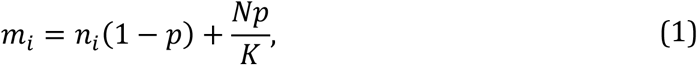

which is the number of individuals that remained in the *i*-th patch plus the number of individuals that arrive to the *i*-th patch after the switch.

After the switch, all individuals will remain in their new food patch until the instantaneous feeding rate reaches *g*_*D*_. This will happen at time 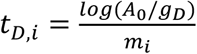 for the i-th food patch. It’s now convenient to split the calculation of the total intake (*G*) in the two periods before and after dispersal. Then, for an individual that spends its time between the switch and the dispersal at the *i*-th patch, the total intake is

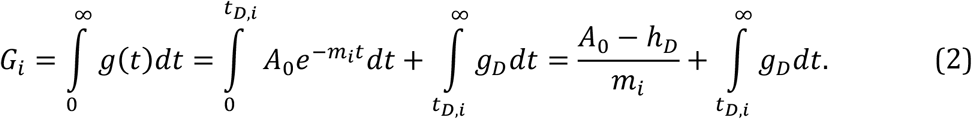

Note that the solution to the first integral is just the amount of food eaten in the patch (*A*_0_ − *g*_*D*_), distributed equally across the *m*_*i*_ individuals populating it.

Now we can compute the expected payoff for each decision (〈*H*〉), which is the expected total food intake minus any costs incurred by the behavior. Individuals that switch have an equal probability of ending up in any of the food patches, so their expected payoff is simply the average of the payoffs across the food patches minus the cost of switching:

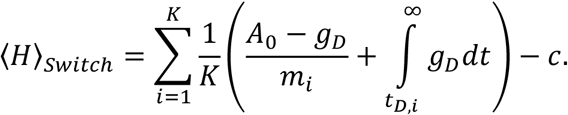

In contrast, individuals that remain have a probability *n*_*i*_/*N* of being in the *i*-th patch, so their expected payoff is

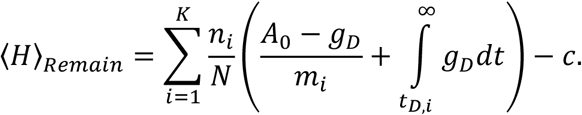

We now compute the benefit of switching,

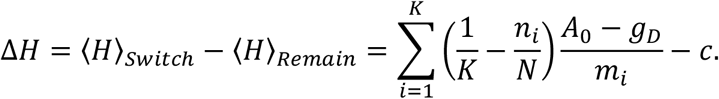

It is now convenient to define

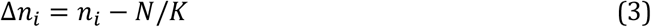

which is the deviation in the initial number of individuals from the average number of individuals in every food patch. We also substitute *m*_*i*_ according to Equation 1, getting

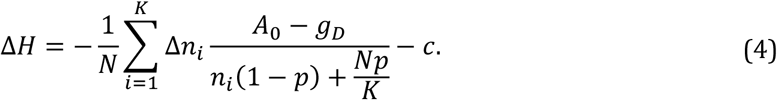

The equilibrium value of *p*, or *p*\* will be such that Δ*H* = 0, so

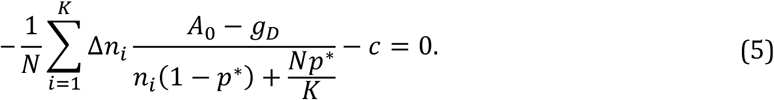

We did not find a simple analytical expression for the value of *p*\*, but we can make several observations:

- If all Δ*n*_*i*_ are zero, the first term of Δ*H* is always zero, so Δ*H* ≤ 0 and switching is never advantageous (therefore, *p*\* = 0). This makes intuitive sense: If all Δ*n*_*i*_ are zero, the individuals were initially distributed in the optimal way (equally distributed across the food patches), so switching cannot bring any benefit.
- If *c* = 0, then *p*\* = 1 regardless of the value of the rest of the parameters (this can be checked by substitution in Equation **Error! Reference source not found.**). This recovers the result of our costless model, in which all individuals should switch regardless of the values of the other parameters.
- When *p* = 1, Δ*H* = −*c*
- When *p* = 0, 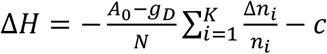
- Δ*H* decreases monotonically as *p* increases, for any values of the parameters.^3^

Therefore, Δ*H* decreases monotonically between 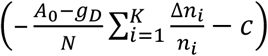 and − *c*, and the value of *p* at which Δ*H* = 0 the equilibrium *p*\* of our Evolutionary Stable Strategy (**Figure S3**). If 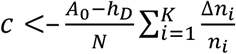, then *p*\* is greater than 0 and a fraction of the population will switch. If 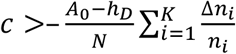, then *p*\* = 0.

In the equilibrium, no mutant has an incentive to deviate from its strategy, since both switching and remaining give the same payoff. Furthermore, the equilibrium is stable: When *p* > *p*\*, Δ*H* becomes negative, meaning that individuals that switched get lower payoff than individuals that remained, and pushing the population towards lower *p*. Conversely, when *p* < *p*\*, individuals that switched have an advantage and the system is pushed towards higher *p*.

##### 3. Individuals must not switch more than once

Individuals that switched at have the same probability of being in every food patch. A new switch will leave these probabilities unchanged^4^, so will not affect the expected payoff. Therefore, individuals have no incentive to switch more than once.

##### 4. Individuals must switch at *t* = 0

A mutant that delays the switch to some later time *t*_*S*_ > 0 will spend its time before the switch in an overcrowded patch (on average) and will therefore get lower final payoff than the wild-type that switches at time *t* = 0. Let’s see it mathematically, comparing the expected payoff for switching at *t* = 0 and the expected payoff for switching at *t* = *t*_*S*_:

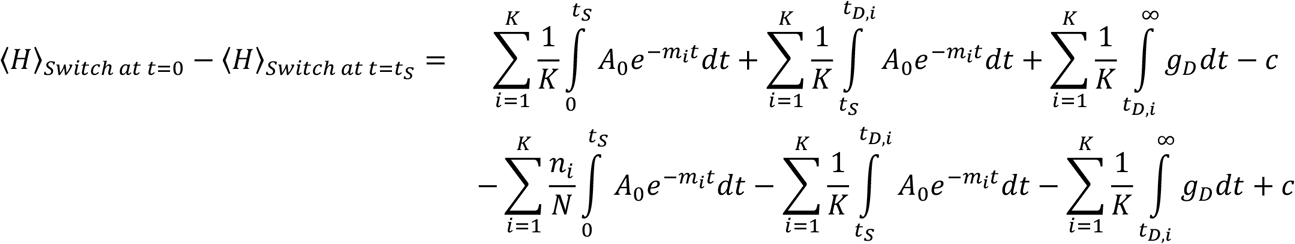

Most of the term cancel out, leaving

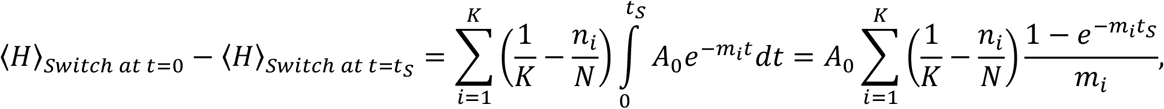

which is always positive^5^. Therefore, switching at *t* = 0 is advantageous.

##### 5. Individuals must disperse once their current food patch is depleted

Once *g*(*t*) = *g*_*D*_, remaining in the same patch will lead to an instantaneous feeding rate below *g*_*D*_, because *g*(*t*) decreases over time. Therefore, worms should not remain. And neither they should switch, as we saw in sections 3 and 4. Therefore, they should disperse.

#### Discussion

A key assumption of the model is that the food patches give diminishing returns, meaning that for individuals remaining in a food patch *h*(*t*) decreases over time. In some cases, this may not be true, for example if food density is so high at the beginning that animals are feeding at their maximum possible rate until food depletion reaches some threshold. In this case, switching is not advantageous until this threshold is met.

**Figure S3.**
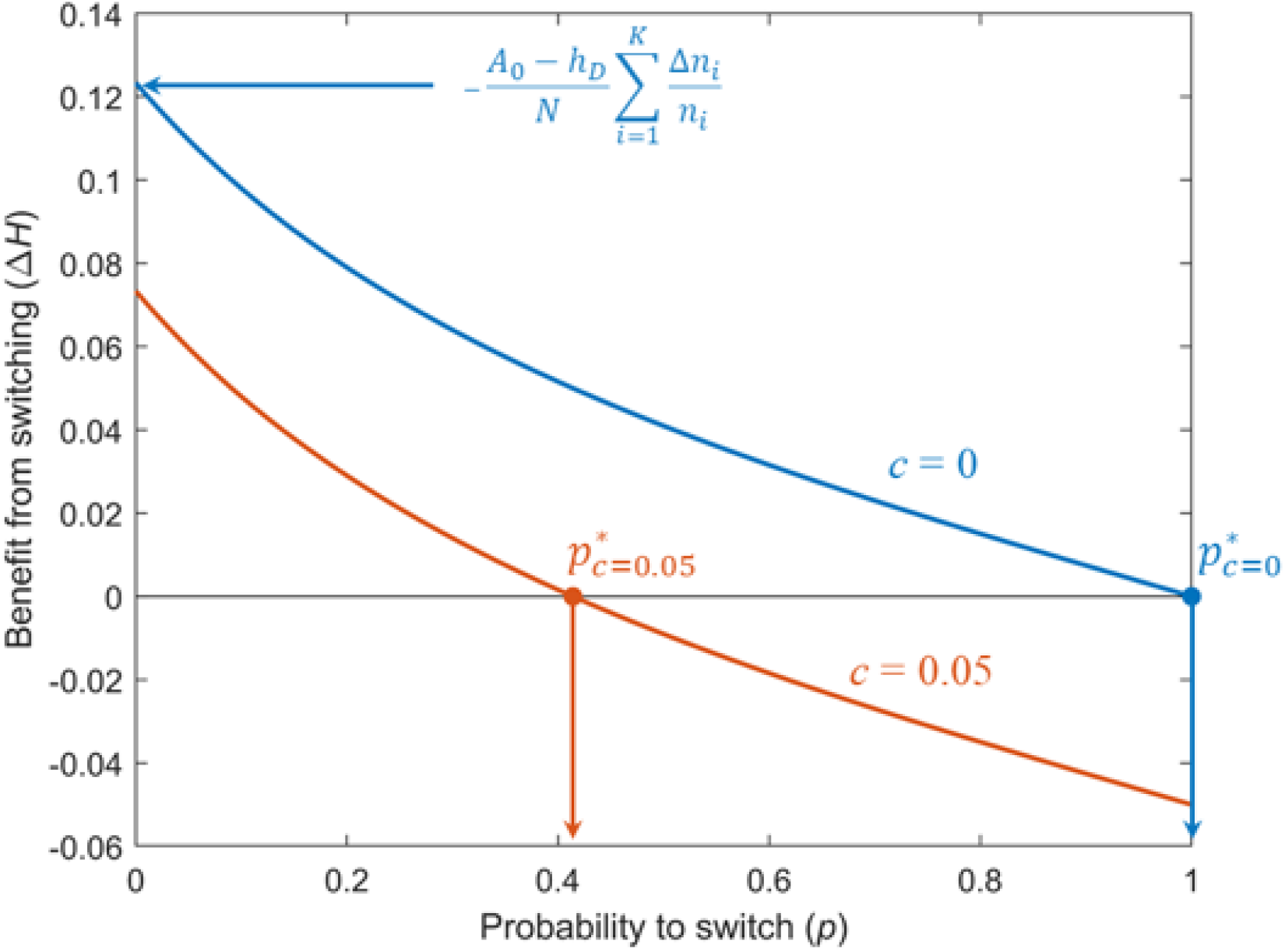
Benefit that individuals obtain from switching at time *t* = 0, with respect to individuals that remain (Equation **Error! Reference source not found.**), with respect to the probability of switching. Blue: Cost *c* = 0. Orange: Cost *c* = 0.05. For both curves, *K* = 5, *n*_*i*_ = *i*, *N* = 15, *h*_*D*_ = 0.1 and *A*_0_ = 1.1.

1 In general, we have *g*(*t*) = *Ca*_*i*_(*t*), where *C* is a constant. But we simply amounts to re-scaling the units of time).

2 2 Proof: If *m*_*i*_ worms occupy a patch, and each worm feeds at a rat a rate 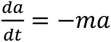. Assuming that mi remains constant over time, the *a*(*t*) = *A*_0_*e*^−*mt*^, where *A*_0_ is the initial food density.

3 Proof: 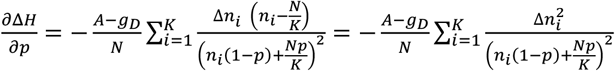. This is always negative as long as *A* > *g*_*D*_, because *N* is always positive and all terms inside the sum are squared.

4 We assume that the population is large enough so that a single mutant does not alter the distributions significantly.

5 **Proof:** 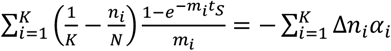, where Δ*n*_*i*_ is defined as in Equation (3), and we define 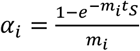. We will show first that Δ*n*_*i*_ and *α*_*i*_ are perfectly anticorrelated (i.e. if Δ*n*_*i*_ > Δ*n*_*j*_, then *α*_*i*_ < *α*_*j*_ for any *i*, *j*). Then, we will show that this implies that 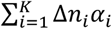 must be negative. **Δ*n*_*i*_ and *α*_*i*_ are perfectly anticorrelated:** Both Δ*n*_*i*_ and *α*_*i*_ depend on *n*_*i*_. Let’s see that their derivatives with respect to it have opposite signs: From Equation (3), 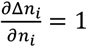, so it’s positive. From Equation (1), 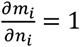, so 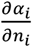 has the same sign as 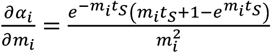. Given that 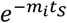 and 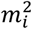 are always positive, this has the same sign as 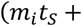 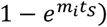, which is always negative (it’s zero for *m*_*i*_ = 0, and 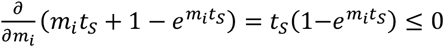 as long as *t*_*S*_ ≥ 0 and *m*_*i*_ ≥ 0, which is always true. Therefore, 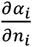 is always negative. 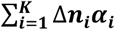 **is always negative:** Let’s split the sum, separating the terms according to the sign of Δ*n*_*i*_: 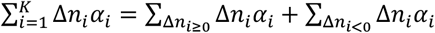. Now, given that all the Δ*n*_*i*_ in the first sum are greater than all the Δ*n*_*i*_ in the second, and that Δ*n*_*i*_ and *α*_*i*_ are perfectly anticorrelated, all the *α*_*i*_ in the first sum must be smaller than all the *α*_*i*_ in the second. Therefore, we can find a number *α*_0_ which is in the middle of the two groups of *α*_*i*_, so that 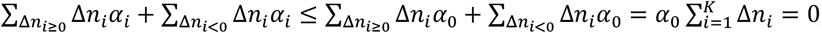,

